# A gradient from long-term memory to novel cognition: transitions through default mode and executive cortex

**DOI:** 10.1101/2020.01.16.908327

**Authors:** Xiuyi Wang, Daniel S. Margulies, Jonathan Smallwood, Elizabeth Jefferies

**Affiliations:** Department of Psychology, University of York, Heslington, York, YO10 5DD, United Kingdom; Centre National de la Recherche Scientifique (CNRS) UMR 7225, Frontlab, Institut du Cerveau et de la Moelle Épinière, Paris, France

**Keywords:** semantic, gradient, default mode network, semantic control, multiple demand

## Abstract

Human cognition flexibly guides decision-making in familiar and novel situations. Although these decisions are often treated as dichotomous, in reality, situations are neither completely familiar, nor entirely new. Contemporary accounts of brain organization suggest that neural function is organized along a gradient from unimodal regions of sensorimotor cortex, through executive regions to transmodal default network. We examined whether this graded view of neural organization helps to explain how decision-making changes across situations that vary in their alignment with long-term knowledge. Functional magnetic resonance imaging found that as decisions are made in an increasingly familiar context, the BOLD signal follows this neural gradient, with stronger responses in default regions when items are linked in long-term memory. In this way, neural organization is optimized to support decision-making in both highly familiar and less familiar situations.

## Introduction

Cognition supports behavioral flexibility by guiding decision-making in a manner that is appropriate to our changing circumstances. Although long-term knowledge can support our decisions in familiar situations, in novel circumstances, knowledge is not available to constrain our choices^1, 2^. Traditionally, these different facets of cognition have been ascribed to dichotomous neural systems. Regions within the lateral anterior temporal lobe and angular gyrus, allied to the default mode network (DMN), are assumed to play a role when the knowledge required by a task is readily available within long-term memory^3–10^. In the absence of long-term knowledge, decisions are supported by the multiple demand network (MDN), including the inferior frontal sulcus, intraparietal sulcus, and pre-supplementary motor area, which support short-term goals in working memory^11, 12^.

Although familiar and novel situations are often treated as dichotomous, many everyday contexts are neither completely familiar, nor entirely novel. Contemporary accounts of cortical organization suggest that neural organization is organized along a gradient that extends from primary sensorimotor areas at one end, through attention and executive areas of MDN, to transmodal DMN regions at the opposite end^13^. This organizational property of the cortex is reflected in its geometry, with the default mode network located in regions of cortex that are most distant from sensory-motor systems. Since there are multiple DMN peaks in the brain, there are multiple spatial gradients (or ‘zones’) extending along the cortical surface from the DMN – with similar patterns within temporal, medial and lateral prefrontal cortex ^13–18^. In this way, the macroscale organization of the cortex highlighted by the principal connectivity gradient offers a view of neural organization in which the relationships between systems are graded rather than dichotomous, with transitions between networks in multiple cortical zones following the same orderly sequence.

Motivated by this novel view of how neural function is organized, the current study examines whether this cortical gradient^13^ reflects how decision-making changes in a graded fashion as the support from long-term knowledge is varied. We capitalized on the fact that semantic long-term memory captures shared features between similar items drawn from the same taxonomic category. We created a novel experimental paradigm in which participants decided whether concepts shared a specific feature (such as color), whilst varying their overall semantic similarity. Participants were asked to decide if two words, presented successively, shared one specific feature (colour, shape or size). We varied the number of other features that the two items shared parametrically, creating a spectrum of trials ranging from situations in which only the goal-relevant feature was shared (e.g., colour: tomato and stop sign) to trials in which nearly all features were shared (e.g., colour: cherry and strawberry).

We measured neural activity in thirty participants using functional magnetic resonance imaging (fMRI) to establish the correspondence between the connectivity gradient described by Margulies et al.^13^ and our ‘task gradient’ – i.e., changes in the neural response to the feature-matching task as the semantic similarity varied. Trials with more shared features are expected to generate stronger responses in regions of the DMN, while concepts that are less globally-overlapping are expected to produce stronger responses in MDN and sensorimotor regions. In order to be able to independently describe DMN and MDN, we used well-established paradigms that manipulated cognition in an independent session.

## Results

### Semantic feature matching performance

Participants saw two words in succession and judged whether they shared a specific feature (colour, shape or size; Figure 1A). We parametrically varied the global semantic similarity of these two concepts (Figure 1B), while eliminating concurrent variation with psycholinguistic variables (see Supplementary Materials). Global feature similarity ratings for matching trials correlated with both accuracy (r = 0.35, p = 0.000; Figure 1C) and response times (r = −0.22, p = 0.007; Figure 1D), indicating that participants could more readily judge that items matched on the current goal feature when task-irrelevant characteristics were also shared. One third of the trials were ‘no match’ trials, and these also varied from globally semantically-related to unrelated. For these decisions, global similarity hindered rather than aided the decision, leading to poorer accuracy when many features were shared (Supplementary Figure S2 shows the behavioural results for non-matching trials).

**Figure 1.**
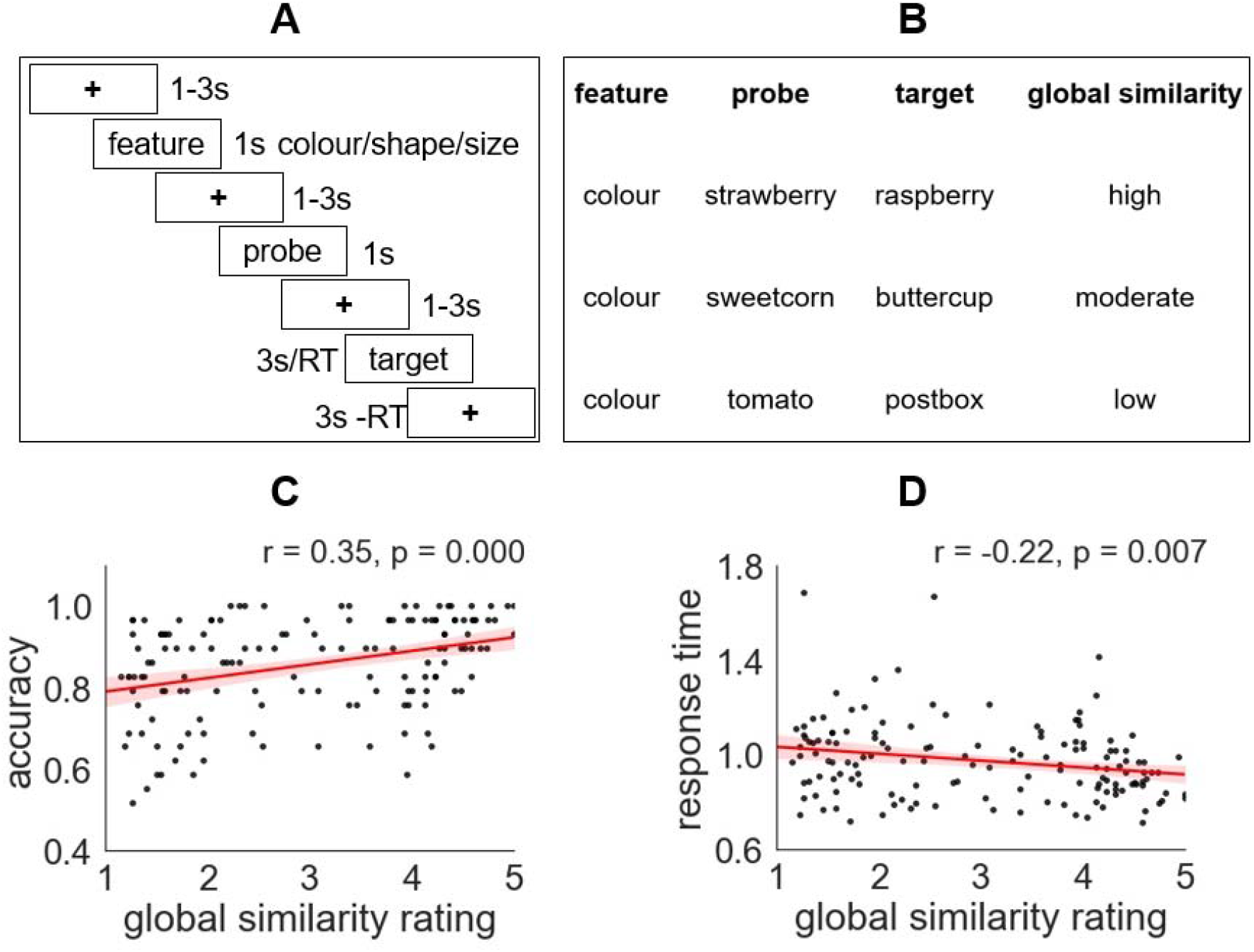
A – Task structure: first, the feature to be matched was specified (e.g. colour), then the probe word was presented (e.g. strawberry), followed by the target word (e.g. raspberry). Participants indicated if the probe and target shared the specified feature. B – Parametric manipulation of global semantic similarity of matching trials within a semantic feature matching task. We created a ‘task gradient’ varying from strong to weak alignment between the inputs and long-term memory. C and D show correlations between ratings of global semantic similarity and average performance across 30 participants (each trial is shown as a data point).

### The parametric effect of global semantic overlap

In order to characterise whole-brain spatial patterns relating to the extent to which the presented words were aligned with the structure of long-term semantic knowledge (i.e. the task gradient), we constructed a model that examined the parametric effect of global feature similarity. In this model, demeaned global semantic similarity ratings were included as a parametric regressor. Figure 2A shows the estimated effect of the parametric manipulation of global feature overlap across the whole brain (i.e., an unthresholded map) for matching trials, when the task-relevant feature was shared between probe and target (supplementary Figure S4 shows the main effect of the task; Figure S4 shows the same analysis for non-matching trials in which the task-relevant feature did not match across the probe and target). Positive effects of this variable (i.e., a stronger BOLD response when items share more features) are seen within lateral anterior-to-mid temporal cortex, angular gyrus and medial and superior frontal regions – regions associated with DMN^19^. Negative effects of this variable (i.e., a stronger BOLD response when items share few features) occur in temporal-occipital cortex, intraparietal sulcus, inferior frontal sulcus and pre-supplementary motor area – regions that fall largely in MDN^11, 12^ (see below for network analysis).

**Figure 2.**
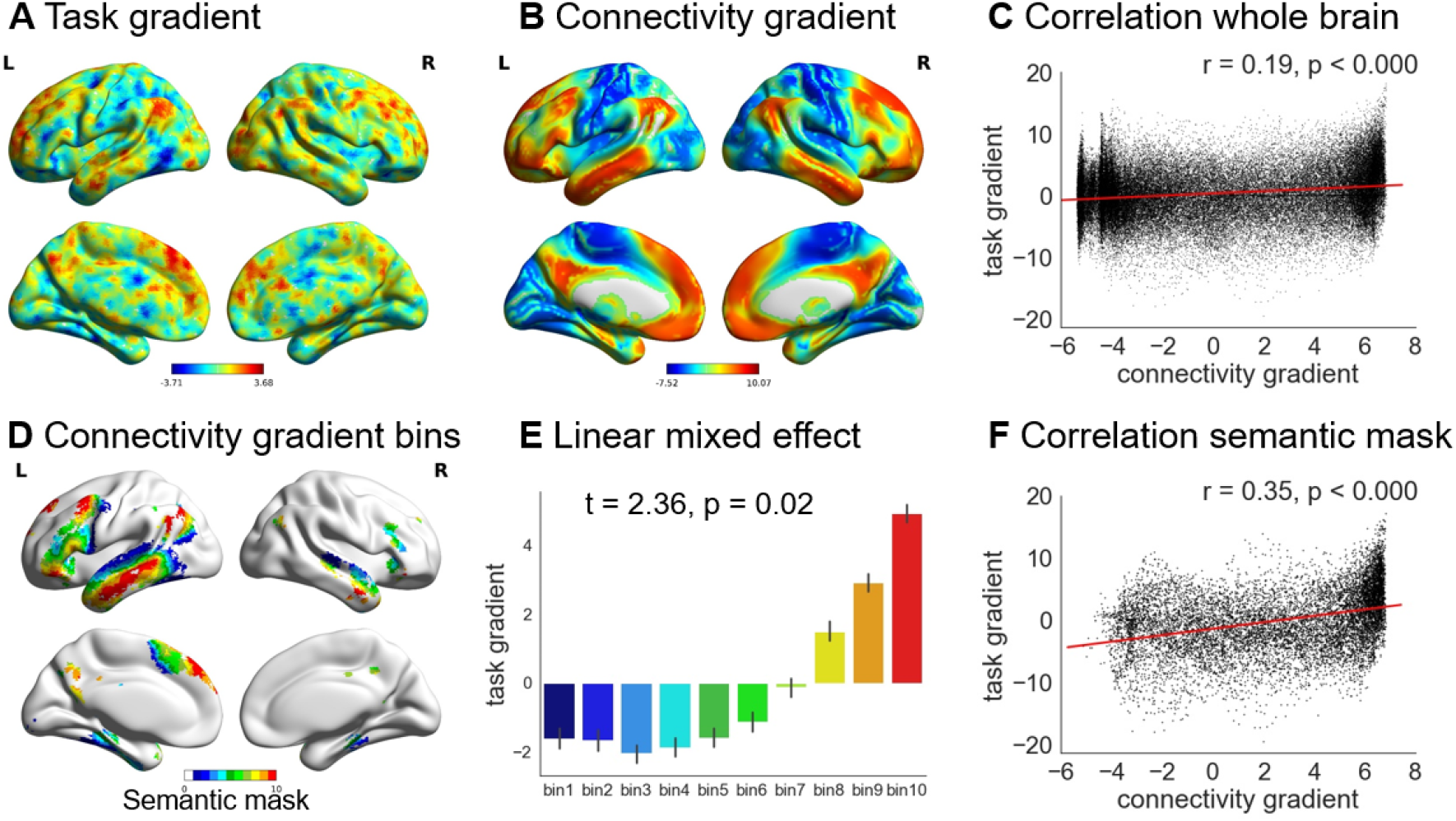
A – Unthresholded map of the task gradient: i.e. parametric manipulation of global semantic similarity, for trials in which the task-relevant feature was shared across probe and target. Warm colours = positively correlated activity [stronger response when both task relevant and task-irrelevant semantic features were shared between probe and target]. Cool colours = negatively correlated activity [stronger response when only the goal feature was shared between probe and target]. B – The principal gradient of intrinsic connectivity from Margulies et al.^13^. C – Correlation between the task gradient for matching trials and the connectivity gradient^13^ across the whole brain. D – The connectivity gradient ^13^ within a semantic mask defined using Neurosynth, divided into decile bins according to gradient value: Bin 1 is located towards the unimodal end, while bin 10 is at the heteromodal end of the principal gradient. E – The effect of the task gradient in each bin of the connectivity gradient within the semantic mask, showing that the response to the task changes in an orderly way. F – Correlation between the task gradient for matching trials and the connectivity gradient^13^ within the semantic mask defined using Neurosynth.

### Correlation between the task gradient for matching trials and the connectivity gradient

Margulies and colleagues found the principal gradient of connectivity (Figure 2B) was anchored at one end by heteromodal DMN regions, and at the other end by unimodal sensory-motor cortex ^13^. We found a significant positive correlation between this connectivity gradient and the task gradient (corresponding to the effect of global semantic overlap when the probe and target shared the task-relevant feature). This correlation was significant across the whole brain (Figure 2C) and was stronger when only voxels associated with semantic processing (falling within a semantic mask from Neurosynth) (Figure 2D) were included (Figure 2F). There was a significant difference between these two correlation coefficients (z = 26.48, p < 0.001), showing that the relationship between the task gradient and the connectivity gradient is to some extent specific to brain regions relevant to semantic processing. A similar positive correlation between the connectivity gradient and the task gradient was observed for non-matching trials, when the target feature was not shared between the probe and target concepts (Supplementary Figure S4), even though global semantic similarity had the opposite effect on behavioural performance for matching and non-matching trials. This suggests that task difficulty is unlikely to fully account for systematic change in functional recruitment along the connectivity gradient.

Next, to test whether a similar functional organisation was present in multiple cortical zones, we calculated the correlation between the connectivity gradient^13^ and the task gradient for the matching trials created through our parametric manipulation of global feature overlap. We focused on four anatomically-defined regions: left lateral frontal, left medial frontal, left lateral temporal, and left lateral parietal cortex (see Supplementary Methods), since these sites are extended across the connectivity gradient and they are also broadly implicated in semantic processing^13, 14^. We found a significant correlation between the task gradient and the connectivity gradient in all four zones (Figure 3).

**Figure 3.**
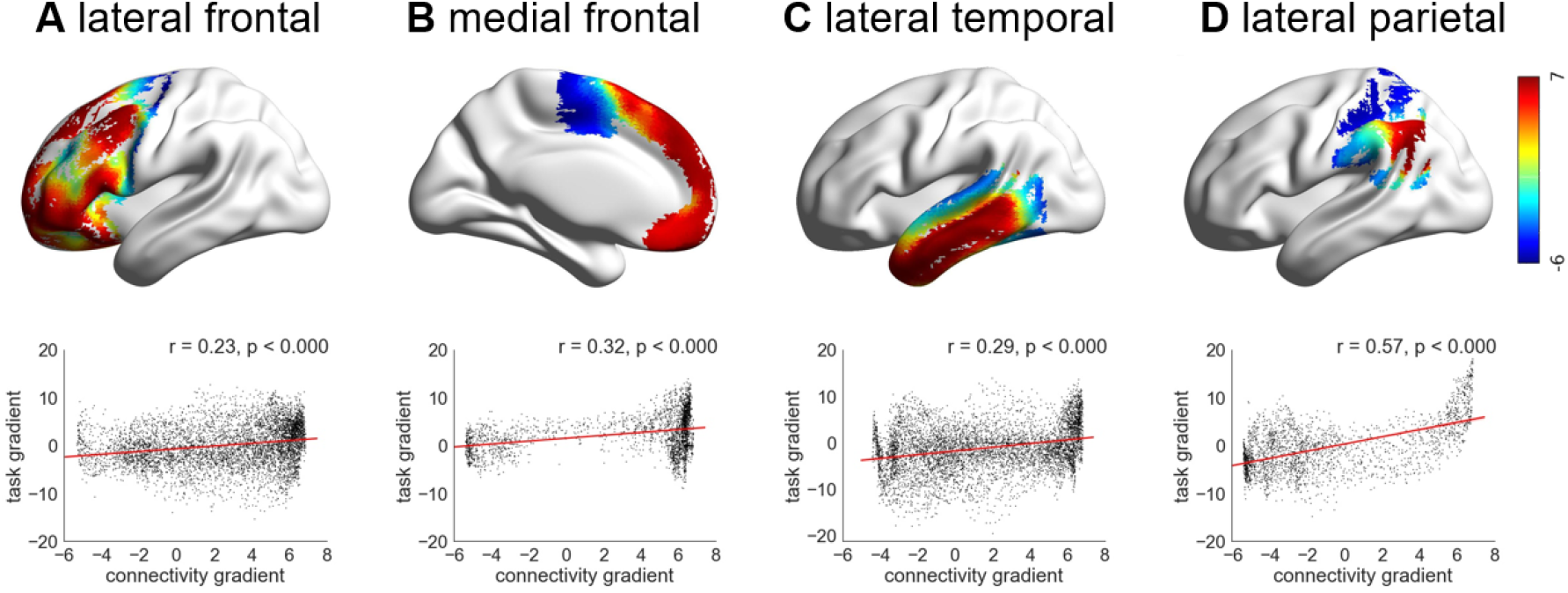
A, B, C and D show the correlation between task gradient for matching trials and connectivity gradient^13^ in lateral frontal, medial frontal, lateral temporal and lateral parietal cortex, respectively. The colour represents each voxel’s value on the connectivity gradient defined by Margulies et al.^13^.

### Systematic change in the effect of the task gradient along the connectivity gradient

To establish if the response to the task gradient for matching trials changed in a systematic way along the connectivity gradient^13^ within the semantic mask, we extracted the beta values corresponding to the effect of global semantic similarity within successive bins along the connectivity gradient (Figure 2D) and examined whether the task response showed a linear change across these bins (Figure 2E). Since the connectivity gradient varies with physical distance from the DMN^13^, a linear trend would suggest that the response to the semantic task changes systematically along the cortical surface. There were ten bins based on deciles, which contained voxels falling within 10% bands along the connectivity gradient, from bin 1 located towards the sensory-motor end of the gradient, through to bin 10 at the heteromodal end overlapping with the DMN (shown in Figure 2D). We characterised the effect of the task gradient within each bin, for each run and for each participant separately, and performed a linear contrast analysis within a linear mixed effects model, including participant and run as a random effect. There was a linear change along the connectivity gradient in the effect of global feature similarity, indicating an orderly relationship between the task gradient and the connectivity gradient^13^. Positive parameter estimates, corresponding to a stronger BOLD response for trials with high global feature similarity in the DMN, gradually reduced in magnitude and became negative at the sensorimotor end of the gradient, reflecting a stronger response for trials with lower global feature similarity (t = 2.36, p = 0.02, Figure 2E). None of the higher order effects, such as quadratic and cubic effects, improved the model fit over the linear effect at p < 0.05. To test the robustness of the linear effect between the effect of the task manipulation and the connectivity gradient bin, we repeated the analysis using 20 bins along the connectivity gradient and observed the same effects (t = 2.72, p = 0.01; Supplementary Figure S7). The parameter estimates and further information about the modelling approach are provided in Supplementary Materials.

### Individuals show the association between the connectivity gradient and task gradient for matching trials

Individual participants are known to show differences in cortical organisation which can be obscured at the group level^20–23^. At the single-subject level, language-selective regions can lie adjacent to multiple-demand regions – and, critically, the location of this functional transition is different across individuals^24^. We therefore examined the voxel-wise correlation between the task gradient for matching trials and the connectivity gradient values^13^ in each individual participant. More than half of the participants showed a significant correlation between the connectivity gradient values and effects of global feature similarity across the whole brain (17/30 participants) at p = 0.05. The same participants showed correlations that exceeded the null distribution based on permutation of the connectivity gradient values^13^. Correlations between connectivity and task gradients were observed in lateral frontal cortex (in 17/30 participants), in medial frontal cortex (in 14/30 participants), in temporal cortex (in 16/30 participants) and in lateral parietal cortex (in 19/30 participants) at p = 0.05. Two-thirds of the sample showed a significant correlation between the task and connectivity gradients within the semantic mask generated using Neurosynth (21/30 participants) at p = 0.05.

### Semantic control peaks are located at the juxtaposition of DMN and MDN

Although prior studies highlight regions of both DMN and MDN as important in semantic cognition, in situations when the support for semantic decisions from long term memory are minimal, the peak neural response is often observed in regions such as the left inferior frontal gyrus, posterior middle temporal gyrus and dorsal anterior cingulate cortex^25^. These peaks typically fall outside the MDN, and are adjacent to, yet distinct from, the DMN^6, 25–27^. Such *semantic control* regions show structural and functional connectivity with regions implicated in long term knowledge (anterior temporal lobes) and domain-general executive control (inferior frontal sulcus)^5^ and our next analysis considers where these regions are located on both the neural gradient observed by Margulies and colleagues, as well as the task gradient generated by our paradigm. Regions within DMN and MDN were defined in the same participants using non-semantic functional localisers. These were contrasts of easy and hard spatial working memory and math judgements taken from Fedorenko et al.^28^. Consistent with previous findings, DMN regions (showing a stronger response to easy versus hard trials in either task) included posterior cingulate cortex, medial prefrontal cortex, angular gyrus and lateral anterior temporal lobes bilaterally. In contrast, MDN regions (showing a stronger response to hard versus easy trials) included inferior frontal sulcus, premotor cortex, intraparietal sulcus, and lateral occipital cortex (Family-Wise Error (FWE)-corrected, z = 3.1, p < 0.05) (Figure 4A).

**Figure 4.**
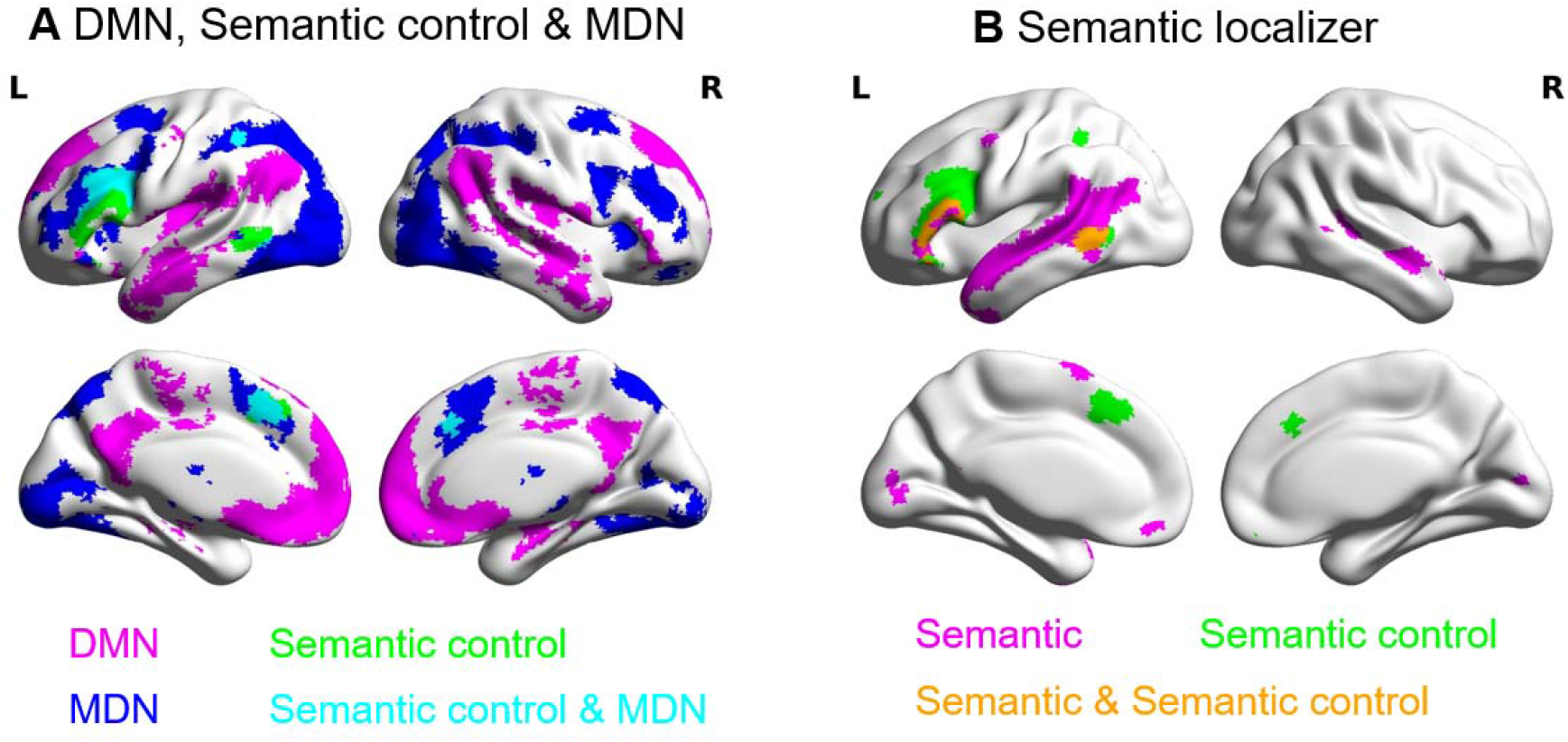
A – Networks used in the analysis, including DMN (in magenta) and MDN (in blue), defined by the localizer tasks, together with semantic control regions from the meta-analysis of semantic control^25^ (in green). B – Semantic regions defined by the semantic localizer task (in magenta), regions implicated in semantic control by an activation likelihood meta-analysis^25^ (in green), and their overlap (in orange). This highlights the critical role of left inferior frontal gyrus and posterior middle temporal gyrus in controlled semantic processing.

Noonan et al.^25^ found left inferior frontal gyrus, left posterior middle temporal gyrus and dorsal anterior cingulate were the most reliably activated sites across different manipulations of semantic control in a neuroimaging meta-analysis. We compared the location of these semantic control sites with DMN and MDN, as defined by the localiser tasks (Figure 4A). The semantic control sites overlapped with MDN in lateral and medial prefrontal regions, in line with the view that MDN is recruited whenever task demands are high. However, two of these sites (left inferior frontal gyrus and middle temporal gyrus) also responded to a semantic localiser, showing stronger activation during the maintenance of strings of words than pronounceable nonwords (FWE-corrected, z = 3.1, p < 0.05) (Figure 4B). The word condition was easier than the nonword condition (see Supplementary Materials) and consequently these semantic control regions did not show the functional profile of the MDN. Instead, in left inferior frontal gyrus, left posterior middle temporal gyrus and dorsal anterior cingulate, the response to semantic control demands, defined by^25^, was observed at the juxtaposition of DMN and MDN. This observation was particularly striking in posterior middle temporal gyrus (Figure 4A), where the semantic control response was juxtaposed between these two canonical networks.

### The task and connectivity gradients capture the orderly transitions between DMN, semantic control and MDN

To further understand the mapping between the DMN, semantic control and MDN networks, we examined their location along both the task gradient and the connectivity gradient^13^. In other words, we extracted the average values for the voxels falling within each network on the parametric map depicting the effect of varying global semantic similarity during feature matching and on the connectivity gradient map. We were then able to test for significant differences in connectivity and task gradient values between these networks.

We defined the following cortical regions within the Neurosynth semantic mask: (i) regions within DMN, (ii) semantic control regions from the meta-analysis of Noonan et al.^25^ that fell outside MDN, (iii) semantic control regions within MDN and (iv) MDN regions not implicated in semantic control (Figure 5A). MDN and DMN were defined using the functional localisers described above. We found that the task gradient values decreased in an orderly fashion across these networks: voxels within DMN had the highest task gradient values, voxels in the semantic control network had lower values, while voxels that fell only in MDN had the lowest beta values (Figure 5B). One-way ANOVA found significant differences in task gradient values (F(3,10987) = 899.97, p < 0.001). Post hoc tests using Tukey HSD criterion for significance indicated significant differences between every network pair (p < 0.05).

**Figure 5.**
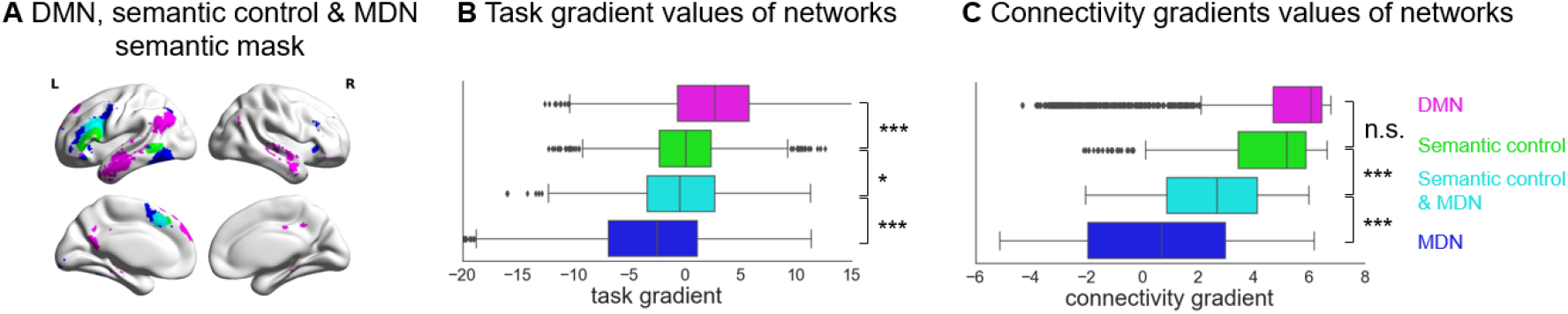
A – Networks used in the analysis. The same networks shown in Figure 4A, including DMN, semantic control and MDN, masked by semantically relevant regions, defined using Neurosynth. B – The task gradient values of each network for matching trials. C – The connectivity gradient values of each network. (* P < 0.05; *** P < 0.001; n.s. = not significant)

We found the same decreasing pattern revealed by the task gradient (Figure 5C). One-way ANOVA found significant differences in connectivity gradient values (F(3, 4990) = 673.57, p < 0.001). Post hoc analysis using the Tukey HSD post hoc criterion for significance indicated significant differences between each pair (p < 0.001) except between DMN and semantic control regions outside MDN (p = 0.309). These findings suggest both the task gradient and connectivity gradient capture the orderly transitions between DMN, semantic control and MDN.

## Discussion

Our study investigated whether contemporary views emphasizing a graded relationship between neural systems can explain how the response to semantic decisions changes systematically across the cortex as the similarity of the presented information with long-term knowledge is varied. We measured neural function using fMRI while participants made decisions about specific conceptual features of words (colour, shape or size), and manipulated the global semantic similarity of the probe and target words parametrically. At one end of this ‘task gradient’, items shared both goal-relevant and irrelevant features, such that these decisions occurred in a context that is supported by long-term memory. At the other end, items had little conceptual overlap beyond those relevant to the goal. Consistent with our hypothesis, we demonstrated that functional recruitment in this task followed the graded organization of the cortex^13^: strong global semantic similarity elicited more activation towards heteromodal DMN regions, while weaker global semantic similarity produced more activation within regions linked to executive control and unimodal processing. The association between our task gradient and the connectivity gradient was observed across multiple points on the cortical mantle, and was most clearly seen in regions associated with semantic cognition. This pattern was observed in individual brains, suggesting that it is not a product of group averaging, and it could not be readily explained in terms of task difficulty, since non-matching trials (in which the probe and target concept did not share the task-relevant feature) showed a similar pattern – even though for these items, global semantic similarity was not associated with a behavioural advantage. Finally, we found that the task gradient acts as an organizing principle for the topographical distribution of neural systems implicated in semantic processing, including the observation that the semantic control network falls mid-way between DMN and MDN.

### The correspondence between the task gradient and the connectivity gradient

Our study suggests that the organisation of the cortex may be optimized to support decision-making across a range of situations that vary from highly familiar to novel. The principal gradient of connectivity is consistent with a hierarchical view of brain organisation, in which heteromodal processing emerges from the gradual integration of unimodal sensory-motor representations. This view has been described previously within the temporal lobes by the ‘graded hub account’ of conceptual representation, which proposes that different modalities (visual, auditory, valence) are gradually integrated within the anterior temporal lobes, with heteromodal conceptual responses falling within ventrolateral regions that are maximally distant from these different inputs^14, 29, 30^. The whole-brain nature of the principal gradient suggests that a similar gradual abstraction of heteromodal representations occurs across the brain^31^. Moreover, the principal gradient of connectivity captures the sequence of large-scale networks on the cortical surface, which show orderly transitions from primary visual/auditory/motor systems, through attention networks, to fronto-parietal control regions, to default mode regions, in multiple zones^13^.

We established this macroscale pattern of functional transitions within a single task when decision-making occurred in contexts that varied in their relevance to long-term memory. When many features of the concepts were overlapping, and consequently the presented information was consistent with how we represent information in long-term memory, trials elicited greater engagement at the heteromodal end of the gradient. Conceptual access in both feature-matching and mismatching tasks would be expected to benefit from overlap in long-term conceptual representations towards the DMN-end of the gradient. In contrast, for trials in which only the goal feature was shared by the two concepts, the functional response was maximal further down the gradient, within attentional networks and unimodal regions. In this way our data highlights one important consequence of the pattern of neural organization described by Margulies et al. (2016)^13^: it may reflect how neural organization is optimized to benefit decisions when the structure of long term memory is variably congruent with newly-presented information.

### Systematic functional transitions in multiple zones

We found correspondence between the task gradient and the connectivity gradient^13^ in multiple cortical zones, including (i) ventral anterior temporal lobe to auditory and visual cortex, (ii) ventromedial to dorsomedial prefrontal cortex, (iii) angular gyrus to intraparietal sulcus, and (iv) inferior frontal gyrus to inferior frontal and precentral sulcus. Although our task involved decisions based on visual features – namely colour, shape and size – functional transitions were observed in regions far from visual cortex. This suggests that the functional gradient is a general principle of whole brain organisation that captures multiple local gradients. Our findings are consistent with several local gradients that have been described in isolation^14, 29–32^. Jackson et al.^15^ identified graded change in the structural and functional connectivity of the ventral medial prefrontal cortex, from DMN to sensory-motor cortex, in line with functional transitions observed in this region^33, 34^. Similarly, Badre et al.^35^ proposed a rostrocaudal gradient in lateral frontal cortex, with rostral frontal areas supporting more abstract forms of control than caudal areas. We observed more complex spatial transitions in lateral frontal cortex, with high gradient values in anterior, ventral and dorsal regions. This is in line with the revised framework of Badre et al.^17^, which suggests that although the frontal lobes are organized hierarchically, there is no unidimensional spatial gradient of abstraction or global difficulty in this region^17, 35, 36^.

### The large-scale task gradient explains the spatial arrangement of networks that support semantic cognition

We found that networks that support semantic cognition are organised in a systematic way along the gradient. Multiple networks with distinct connectivity profiles are recruited in semantic tasks – including DMN regions (such as angular gyrus and lateral middle temporal gyrus) ^37, 38^, brain regions specifically implicated in semantic control^25^, and MDN regions implicated in domain-general executive control^11, 29^; however, the topographical organisation of these networks has not been previously investigated. On both the connectivity and task gradients, the semantic control network had intermediate values – falling in between DMN and MDN in terms of patterns of connectivity and task response. The spatial adjacency of the semantic control network to both DMN and the MDN might allow semantic control regions to integrate long-term conceptual knowledge with more adaptive representations of currently-relevant goals, supporting flexible patterns of semantic retrieval^5^. This is consistent with recent evidence that DMN and MDN cooperate when memory is controlled^39–42^.

Graded transitions in the BOLD response from DMN to executive cortex might reflect a shift from intrinsically-guided retrieval based on representations in memory, to a stronger focus on the specific feature information required by the task, when conceptual combinations do not overlap with long-term memory. Across multiple studies, DMN regions show stronger activation when semantic tasks are guided by memory^7, 9, 38^. In contrast, when current inputs are not a good match with the information in memory, a complementary strategy is needed in which intrinsic cognition is temporarily suppressed. MDN is thought to dynamically alter semantic processing by coding for information relevant to the current decision^12^ and by changing its pattern of connectivity according to task demands^43^. Given MDN activates to demanding tasks while DMN typically deactivates^44^, task difficulty might contribute to the functional gradient that we observed for matching trials. However, on non-matching trials (when the probe and target did not share the task-relevant feature), a similar positive correlation was observed between the task gradient and the connectivity gradient – even though global semantic similarity was no longer associated with easier decisions. This suggests that the functional gradient reflects the semantic overlap between probe and target as opposed to cognitive control demands or task difficulty per se.

Further studies that parametrically manipulate other types of semantic and non-semantic tasks are needed to examine the specificity of our findings. A single study cannot fully specify the critical aspects of the task which gave rise to the functional gradient; for example, this research does not address the question of whether the same spatial relationships would be observed for parametric manipulations of non-semantic goals, and/or other semantic tasks, such as global association tasks, that vary in association strength but in the absence of an explicit goal which requires that participants focus on specific features represented at the unimodal end of the gradient. We found that the correlation between the task gradient and the connectivity gradient was maximal within brain regions recruited during semantic tasks, suggesting that if there are similar patterns for non-semantic tasks, these might be strongest in different cortical regions. The unique contribution of the current study is to show that functional recruitment within a task can show systematic variation in a way that follows graded changes in intrinsic connectivity.

## Materials and Methods

### Participants

The research was approved by local ethics committees and volunteers provided written informed consent. 31 healthy adults were recruited from the University of York (26 females; age: mean ± SD = 20.60 ± 1.68). All participants were right-handed, native English speakers, with normal or corrected-to-normal vision and no history of psychiatric or neurological illness. We excluded one participant who attended only one of two sessions and two participants because accuracy was not higher than chance.

### Design and tasks

Participants matched probe and target concepts (presented as written words) according to colour, shape or size, in a rapid event-related design. The goal feature was specified at the start of each trial. The degree of feature overlap between the probe and target was parametrically manipulated: in some trials, the items shared many features (e.g., `STRAWBERRY` and `RASPBERRY`), while others only shared the task-relevant feature (e.g. colour match: `TOMATO` and `POSTBOX`) (Figure 1). Localiser tasks were presented in a different session to define functional networks of interest. We compared visually-presented words and nonwords to identify sites sensitive to meaning. We also contrasted easy and hard spatial working memory and math decisions to define DMN and MDN^11, 28^. Further details about the tasks are available in the Supplementary Materials.

### MRI data acquisition

Structural and functional data were collected on a Siemens Prisma 3T MRI scanner at the York Neuroimaging Centre. The scanning protocols included a T1-weighted MPRAGE sequence with whole-brain coverage. The structural scan used: acquisition matrix of 176□×□256□×□256 and voxel size 1□×□1□×□1mm_3_, repetition time (TR) = 2300ms, and echo time (TE) = 2.26ms. Functional data were acquired using an EPI sequence with a 80° flip angle and using GRAPPA with an acceleration factor of 2 in 3 x 3 x 4-mm voxels in 64-axial slices. The functional scan used: 55 3-mm-thick slices acquired in an interleaved order (with 33% distance factor), TR = 3000ms, TE = 15ms, FoV = 192mm.

### MRI data pre-processing

Pre-processing was carried out using FMRIB’s Software Library (FSL version 6, fsl.fmrib.ox.ac.uk/fsl/fslwiki/FEAT/). The T1-weighted structural brain images were extracted. Structural images were registered to the MNI-152 template using FMRIB’s linear image registration tool (FLIRT). fMRI data pre-processing included motion correction, slice-timing correction, spatial smoothing with a 5mm FWHM Gaussian filter and high-pass filtering at 100s. Motion-affected volumes were removed from the fMRI data using scrubbing^45^.

## MRI analysis

### Parametric task gradient analysis

We modelled the parametric effect of global feature similarity by entering the demeaned global semantic similarity ratings for correct matching trials as a parametric regressor. We also modelled the main effect of task, the instruction period, two inter-stimulus interval periods, probe presentation, mismatch trials in which probe and target did not share the specified feature, and incorrect trials.

### Correlations between task and connectivity gradient at the group level

We examined the spatial correspondence of the task gradient for matching trials and connectivity gradient by computing correlations across the whole brain and within semantic regions defined using a meta-analytic mask for the term “semantic” from Neurosynth^46^. To establish whether systematic functional change along the connectivity gradient was seen in multiple cortical areas linked to semantic processing, we also examined the correlations between the task and connectivity gradients using anatomical masks in lateral frontal, medial frontal, temporal and parietal areas (details in Supplementary Materials).

### Mixed effects modelling

To examine the shape of gradient transitions within the semantic mask from Neurosynth^46^, voxels were assigned into ten decile bins according to their values on the connectivity gradient^13^. We investigated whether there was a linear (or more complex) relationship between the effect of the task manipulation and the connectivity gradient bin within a linear mixed effect model. Parameter estimates for the effect of global feature similarity were extracted from each bin, for each subject and in each run separately, with the model capturing the hierarchical structure of this data. To allow for individual differences in the effect of the gradient and in the overall BOLD response, we allowed for random intercepts and slopes within the model. We compared a model that only included a linear effect of gradient bin with more complex models, such as models that additionally included quadratic and cubic effects. Further details including fit statistics and parameter estimates are provided in Supplementary Materials.

### Correlations between gradient values and effects of semantic similarity within individuals

We extracted the parametric effect of the task gradient for matching trials in each voxel and each individual participant to establish how many participants showed a significant correlation between the task gradient and the connectivity gradient. We tested these correlations in each participant within the semantic mask and for each anatomical region of interest. We also permuted the connectivity gradient values (10000 iterations) to examine whether the correlation coefficients were significantly greater than zero.

### Localizer task analysis

For the semantic localiser task, we examined the contrast of words with pronounceable nonword task blocks. In the spatial working memory task and math task, we examined the contrast of hard versus easy conditions to define MDN regions, and the reverse to define DMN regions. A grey matter mask was imposed for all analyses and the resulting clusters were corrected for multiple comparisons using FWE detection at a threshold of z > 3.1, p < 0.05. The whole-brain analysis of the localizer tasks allowed us to examine whether semantic control recruits cortical regions that fall between DMN and MDN. Next, to examine whether the task gradient and connectivity gradient captured the organization of semantic networks for matching trials, we extracted these task and connectivity gradient values within DMN, semantic control and MDN regions, and tested whether there was an orderly change in these values between networks.

### Control analysis examining non-matching trials

To examine whether the effects we found for the matching trials were explained by task difficulty, we assessed the parametric effect of global semantic similarity for the non-matching trials (see supplementary materials). We modelled the task gradient by entering the demeaned global semantic similarity ratings for correct non-matching trials as a parametric regressor. For these non-matching trials, global semantic similarity was negatively correlated with accuracy, in contrast to matching trials which showed a positive correlation between global semantic similarity and behavioural performance. Opposite task gradients for matching and non-matching trials would be consistent with these gradients reflecting task difficulty. In contrast, parallel task gradients across matching and non-matching trials cannot be fully explained by task difficulty and might reflect long-term semantic similarity of the inputs. Correlations between the task gradient and connectivity gradient for non-matching trials, together with mixed effects models characterising this relationship, are provided in supplementary materials.

## Data and code availability

All summary data, materials and code used in the analysis are accessible in the Open Science Framework at https://osf.io/pa84c/. Raw fMRI data is restricted in accordance with ERC and EU regulations.

## Author Contributions

X.W., D.S.M., J.S. and E.J. designed research; X.W. performed research; X.W. analyzed data; X.W. D.S.M., J.S. and E.J. wrote the paper.

## Acknowledgments

This study was supported by the European Research Council (Project ID: 771863 - FLEXSEM and Project ID: 646927 - WANDERINGMINDS). We are grateful to Evelina Fedorenko for providing us with the localizer tasks and to Evgeny Gluzman for piloting a preliminary version of the task. We thank Deniz Vatansever and Theodoros Karapanagiotidis for suggestions about data analysis.

## Competing interests

The author declare no competing interests.

## 1 Tasks

### 1.1 Semantic feature matching task

Participants were asked to match probe and target concepts (presented as words) according to a particular semantic feature (colour, shape or size), specified at the start of each trial. We parametrically manipulated the global feature overlap between the probe and target concepts, creating a “task gradient”, using semantic similarity ratings taken from a separate group of 30 participants on a 5-point Likert Scale. For example, in colour-matching, `STRAWBERRY` – `RSAPBERRY` share many features (not just colour), while `TOMATO` – `POSTBOX` share few features besides colour. The global similarity ratings of the selected trials were evenly distributed from 1 to 5. This parametric design allowed us to model the effect of global feature similarity in the neural data, and test for systematic changes in cortical recruitment along the connectivity gradient defined by Margulies et al.^1^, as semantic similarity increased. Participants were also required to detect non-matches (presented on one third of the trials), which could either be globally-related or unrelated. For example, `LEMON` and `LIME` share many features but not colour, while `BANANA` and `RUBY` share almost no features, including colour. This ensured that participants had to pay attention to the target feature on each trial. Participants pressed a button with their right index finger to indicate a match trial, and responded with their right middle finger to indicate a non-match trial. We then computed behavioural efficiency scores across participants for each item, combining response time (RT) with accuracy: i.e., the mean RT for correct responses across participants was divided by the proportion of correct responses for that item^2^. We reversed this measure so that higher efficiency scores corresponded to better performance^3, 4^.

For the matching trials, global feature similarity ratings positively correlated with behavioural performance (accuracy: r = 0.35, p = 0.000, Figure S2A; response time: r = −0.22, p = 0.007, Figure S2B; efficiency: r = 0.31, p = 0.000, Figure S2C), indicating that participants could more readily identify matching features when task-irrelevant characteristics were also shared between the probe and target concepts. For these matching trials, there were no significant correlations between global semantic similarity and word length (number of letters; r = −0.07, p = 0.429), word frequency (based on SUBTLEX-UK: Subtitle-based word frequencies for British English^5^; r = 0.05, p = 0.586) or word concreteness^6^ (r = 0.07, p = 0.394) of the target word, as shown in Figure S1. There was a weak and non-significant correlation between behavioural efficiency and word frequency (r = 0.14, p = 0.092). There were significant correlations between behavioural efficiency and word concreteness (r = 0.24, p = 0.007) and word length (r = −0.19, p = 0.024). However, semantic similarity ratings correlated positively with behavioural efficiency after regressing out word length, word frequency and concreteness (r = 0.33, p < 0.000), indicating that higher global feature similarity was associated with more efficient feature matching.

Our main analyses concerned these matching trials, for which we had twice as many observations and a smooth spread of trials across all levels of global semantic similarity (see results in main text). However, global semantic similarity had differing effects on behavioural performance in matching and non-matching trials. Consequently, the comparison of these trial types can establish whether functional change along the task gradient is likely to reflect the global semantic overlap between probe and target words, or alternatively the difficulty of the semantic decision. For non-matching trials, global feature similarity ratings showed significant negative correlations with accuracy (r = −0.33, p = 0.005, Figure S2D) and behaviour efficiency (r = −0.33, p = 0.005, Figure S2F). Non-matching trials with higher global feature similarity ratings were also slower, although the correlation with response time did not reach significance (r = 0.20, p = 0.099, Figure S2E). Therefore, in contrast to matching trials, which were easier when global semantic similarity was higher, if anything, greater global semantic similarity made it more difficult to decide that a target feature did *not* match across probes and targets. This relationship between global similarity and behavioural performance was not explained by psycholinguistic confounds. There were no significant correlations for non-matching trials between global semantic similarity and target word length (number of letters; r = −0.05, p = 0.684, Figure S1D), or concreteness^6^ (r = - 0.02, p = 0.867, Figure S1F), as shown in Figure S1. There was a significant correlation between global semantic similarity and target word frequency (based on SUBTLEX-UK^5^; r = 0.25, p = 0.036; Figure S1E). However, the psycholinguistic variables did not correlate with behavioural efficiency, including word length (r = −0.01, p = 0.919), word frequency (r = −0.02, p = 0.886) and concreteness (r = 0.03, p = 0.809). Furthermore, semantic similarity ratings were significantly negatively correlated with behavioural efficiency after regressing out word length, word frequency and concreteness (r = −0.32, p = 0.011), indicating that higher global feature similarity was associated with less efficient no-match decisions. In summary, if the positive relationship for matching trials between the task gradient and the connectivity gradient (reported in the main text) reflects task difficulty (i.e., greater activation towards the DMN-end of the gradient for easier decisions), we would *not* expect to see the same pattern for non-matching trials. In contrast, if this relationship reflects the global semantic similarity of the probe and target words, a similar pattern may be seen for matching and non-matching trials.

In order to maximize the statistical power of the rapid event-related fMRI data analysis, the stimuli were presented with a temporal jitter, randomized from trial to trial^7^. The inter-stimulus intervals (between instruction and probe, and probe and target) and the inter-trial interval varied from 1 to 3s. Each trial started with a fixation, followed by a task instruction slide specifying the feature to match (colour, shape or size), presented for 1s. This was followed by the second fixation and then the probe word, presented for 1s. Finally, there was the third fixation followed by the target word, triggering the onset of the decision-making period. The target remained visible until the participant responded, or for a maximum of 3s. The instruction, probe and target words were presented centrally on the screen.

There were 216 trials in total, presented in 4 runs of 54 trials each. Each run lasted for 600s. Global feature similarity was evenly distributed in each run. Within each run, there were 18 trials for each feature. 12 of these 18 trials were matching trials in which probe and target shared the relevant feature, while 6 were non-matching trials. The order of runs and trials within each run was randomized across subjects. The stimuli were presented using Psychopy^8^.

### 1.2 Localizer tasks

Localizer tasks were used to define default mode network (DMN), multiple demand network (MDN) and semantic regions (adapted from 9, 10). These tasks are shown in Figure S3. MDN regions were defined using contrasts of hard over easy spatial working memory or arithmetic judgements, while DMN regions were defined using the reverse contrasts. Each localizer task included two runs and two conditions, presented in a standard block design. Condition order was counterbalanced across runs and run order was counterbalanced across participants for each task.

#### 1.2.1 Spatial working memory task

Participants had to keep track of four or eight sequentially presented locations in a 3×4 grid ^9^, giving rise to easy and hard spatial working memory conditions. Stimuli were presented at the centre of the screen across four steps. Each of these steps lasted for 1s and highlighted one location on the grid in the easy condition, and two locations in the hard condition. This was followed by a decision phase, which showed two grids side by side. One grid contained the locations shown on the previous four steps, while the other contained one or two locations in the wrong place. Participants indicated their recognition of these locations in a two-alternative, forced-choice paradigm via a button press and feedback was immediately provided. Each run consisted 12 experimental blocks (6 blocks per condition and 4 trials in a 32 s block) and 4 fixation blocks (each 16s long), resulting in a total time of 448s.

#### 1.2.2 Math task

Participants saw an arithmetic addition expression on the screen for 1.45s and were then given two numbers as choices. The easy condition included smaller single-digit numbers while the hard condition included larger two-digit numbers, also presented for 1.45s. Each trial ended with a blank screen lasting for 0.1s. Each run consisted of 12 experimental blocks (with 4 trials per block) and 4 fixation blocks, resulting in a total time of 316s.

#### 1.2.3 Semantic localizer task

Participants read sentences and lists of pronounceable nonwords, followed by a probe recognition test (present/absent judgment for single word and nonword). Sentences contrasted with nonwords reliably activate semantic and language regions^9, 11–13^. Each trial started with a 100ms blank screen. Stimuli were presented at the centre of the screen, one word/nonword at a time, at the rate of 450ms per item. The sequence was followed by a probe word/nonword; participants had 2s to decide whether this item had been presented in the sequence, giving a total trial duration of 7.5s. Each run included 16 experimental blocks with 3 trials per block and 5 fixation blocks lasting for 14s. Each run lasted a total of 430s.

Given that the sentences were easier to maintain than the nonwords, the semantic localiser was able to reveal regions involved in semantic processing (including semantically-relevant regions within DMN and potentially also semantic regions responding to control demands outside DMN). However, this localiser contrast is likely to miss MDN regions, which are expected to respond to the difficulty of the nonword condition but might nevertheless contribute to demanding semantic tasks. Consequently, we defined an additional semantic control mask using a meta-analysis of task contrasts in which semantic control demands were manipulated^14^. This semantic control map might be expected to overlap with the semantic localiser in control regions specific to semantic processing outside the MDN, as well as other control regions within the MDN.

## 2. Supplementary details about analysis

### Masks used to establish if there is a relationship between the connectivity gradient and task gradient in multiple brain regions

To establish whether functional change along the connectivity gradient is seen in multiple cortical areas linked to semantic processing, we examined anatomical masks in lateral frontal, medial frontal, lateral temporal, and lateral parietal areas, in addition to whole-brain analyses. We used the Harvard-Oxford cortical structural atlas and removed voxels with < 5% grey matter probability. For the left lateral frontal cortex, we included frontal orbital cortex, frontal pole, inferior frontal gyrus pars opercularis, pars triangularis, middle frontal gyrus, superior frontal gyrus and frontal operculum cortex, excluding medial prefrontal regions beyond x > −16. This value was set based on cortical anatomy to divide the frontal lobe into medial and lateral portions. For the left medial frontal cortex, we included lateral medial cortex, frontal pole, supplementary motor area and superior frontal gyrus and selected voxels where x > −16 to exclude more lateral regions. For the left temporal area, we included temporal pole, anterior, posterior and temporooccipital inferior temporal gyrus, anterior, posterior and temporooccipital middle temporal gyrus, anterior and posterior superior temporal gyrus, anterior and posterior temporal fusiform cortex, and temporal occipital fusiform cortex. For the parietal area, we included angular gyrus, anterior and posterior supramarginal gyrus, and superior parietal lobe, and we selected voxels where z < 10 to exclude voxels with a high probability of falling in the temporal lobe.

### Mixed effects modelling

To investigate the relationship between the effect of the task manipulation and the connectivity gradient bin, we used the package ‘lme4’^15^ in R version 3.6.1 ^16^ to perform mixed effects modelling. Parameter estimates for the task gradient in matching trials (i.e. the effect of global feature similarity) were extracted for 10 decile bins along the connectivity gradient, for each subject and in each run separately. The mixed-effects model included a single fixed effect of connectivity gradient bin along with random intercepts for subjects and runs, as well as by-subject random slopes for the effect of connectivity gradient bin ^17^. We modelled the linear effect of gradient bin as follows: Y = β_0_ + β_1_ * Bin + r. Y is the beta value modulated by the global similarity rating, β_0_ is intercept, β_1_ is the linear rate of change across gradient bins and r is the residual. We examined models that additionally included higher order effects of gradient bin, including quadratic and cubic effects up to the ninth-order effect, to examine whether these additional terms improved the model fit according to −2 log likelihood. The parameter estimates are provided in Tables S1 and S2. Finally, to test the robustness of the linear effect between the effect of the task manipulation and the connectivity gradient bin, we divided all the voxels within the semantic mask into 20 bins according to their functional connectivity gradient values (Figure S7A) and then repeated the linear mixed effects analysis.

### Data analysis for non-matching trials

To establish whether the effect we found for the matching trials could be explained by task difficulty, we examined the non-matching trials, in which probe and target did not share the specified feature. For these items, global semantic similarity was associated with poorer accuracy and no significant change in response time (while for the matching trials, global semantic similarity was linked to better performance). We modelled the parametric effect of global feature similarity by entering the demeaned global semantic similarity ratings for correct non-matching trials as a parametric regressor. We also modelled the main effect of task, the instruction period, two inter-stimulus interval periods, probe presentation, matching trials in which probe and target shared the specified feature, and incorrect trials. We examined the spatial correspondence between the task gradient for non-matching trials and the connectivity gradient by computing correlations across the whole brain and within semantic regions defined using a meta-analytic mask for the term “semantic” from Neurosynth^18^. To establish whether systematic functional change along the connectivity gradient was seen in multiple cortical areas linked to semantic processing, we examined anatomical masks in lateral frontal, medial frontal, temporal and parietal areas. Finally, we investigated whether there was a linear (or more complex) relationship between the effect of global semantic similarity and the connectivity gradient bin for the non-matching trials, using a linear mixed effects model (see above).

## 3. Supplementary Results

### The parametric effect of global semantic overlap and mean effect of the task

Figure S4 shows the mean effect of the semantic task alongside the parametric effect of the task gradient for matching trials and non-matching trials, respectively. The task was shown to elicit significant activation in ‘task-positive’ regions of the MDN, including inferior frontal sulcus, intraparietal sulcus and pre-supplementary motor area (Family-Wise Error (FWE) -corrected, z = 3.1, p < 0.05) for matching trials (Figure S4B) and non-matching trials (Figure S4F).

Figure S4E shows the parametric effect of global feature overlap for the non-matching trials across all voxels (i.e., the unthresholded map of the task gradient). Positive effects of the task gradient (i.e., a stronger BOLD response when items shared more features) can be seen within lateral anterior-to-mid temporal cortex, angular gyrus and medial and superior frontal regions. Negative effects of this variable (i.e. stronger activation for trials with lower global semantic overlap) were found in intraparietal sulcus, inferior frontal sulcus and pre-supplementary motor area. These effects are similar to those observed for the matching trials (Figure S4A). The unthresholded maps characterising the effects of global semantic similarity for matching and non-matching trials were positively correlated (r = 0.11, p < 0.001 across the whole brain; r = 0.20, p < 0.001 within a semantic mask defined by Neurosynth).

### Correlation between connectivity gradient and task gradient for matching versus non-matching trials

In line with the pattern observed for matching trials, there were significant positive correlations between the connectivity gradient and the effect of the task gradient for non-matching trials (Figure S4G, Figure S4H), even though the trials with higher global semantic similarity were no longer easier. Non-matching trials in which the probe and target shared more non-target features elicited more activation towards the DMN-end of the gradient, despite these trials having lower accuracy. We also found a significant correlation between the task gradient and the connectivity gradient in all four anatomical zones both within a semantic mask (Figure S6) and when all voxels within these zones were included (Figure S5; see main text for the equivalent analysis for matching trials). These analyses together suggest that the relationship between the task gradient and the connectivity gradient reported in the main analysis for matching trials did not reflect task difficulty, and instead was associated with the semantic match between the items in long-term memory.

For non-matching trials, the correlations between the task gradient and the connectivity gradient were significantly stronger when including all the voxels in each zone compared with only voxels associated with semantic processing (falling within a semantic mask from Neurosynth), in lateral frontal (z = 7.87, p < 0.001), medial frontal (z = 6.42, p < 0.001) and lateral parietal (z = 2.76, p = 0.006) regions. There was no difference in the strength of this correlation between the connectivity gradient and task gradient for non-matching trials with and without the application of the semantic mask in lateral temporal cortex (z = 0.97, p = 0.33).

For matching trials, the correlations between the task gradient and the connectivity gradient were significantly stronger when only voxels associated with semantic processing (falling within a semantic mask from Neurosynth) were included in the analysis, both in medial frontal (z = 6.16, p < 0.001) and lateral temporal (z = 7.21, p < 0.001) regions. This effect resembles the pattern seen across the whole brain (see main text). There was no difference in the strength of this correlation for matching trials with and without the application of the semantic mask in lateral frontal (z = 0.74, p = 0.46) or lateral parietal cortex (z = 0.53, p = 0.60). Brain maps were visualized using BrainNet Viewer^19^.

### Systematic change along the connectivity gradient

A supplementary linear mixed effects analysis for the matching trials showed an orderly relationship between the task gradient and the connectivity gradient divided into 20 bins, as opposed to 10 in the main analysis (Figure S7A). There was a linear change along the connectivity gradient in the effect of global feature similarity: positive parameter estimates, corresponding to a stronger BOLD response for trials with high global feature similarity in the DMN, gradually reduced in magnitude and became negative at the sensorimotor end of the gradient, reflecting a stronger response for trials with lower global feature similarity (t = 2.43, p = 0.02, Figure S7B). Only the quadratic effect improved model fit over the linear effect (χ2 (1) = 7.95, p = 0.005).^1^ The parameter estimates for this model are provided in Tables S3 and S4.

Figure S7C and S7D show the effect of the task gradient for non-matching trials in each bin of the connectivity gradient within the semantic mask. In the linear mixed effects analysis, the linear effect of global semantic similarity across bins showed a non-significant trend (t = 1.95, p = 0.06) when dividing all the voxels into 10 bins. There was a significant linear effect of global semantic similarity along the connectivity gradient when this was divided into 20 bins (t = 2.12, p = 0.04). The parameter estimates for these two models are provided in Tables S5, S6, S7 and S8.

---

**Figure S1.**
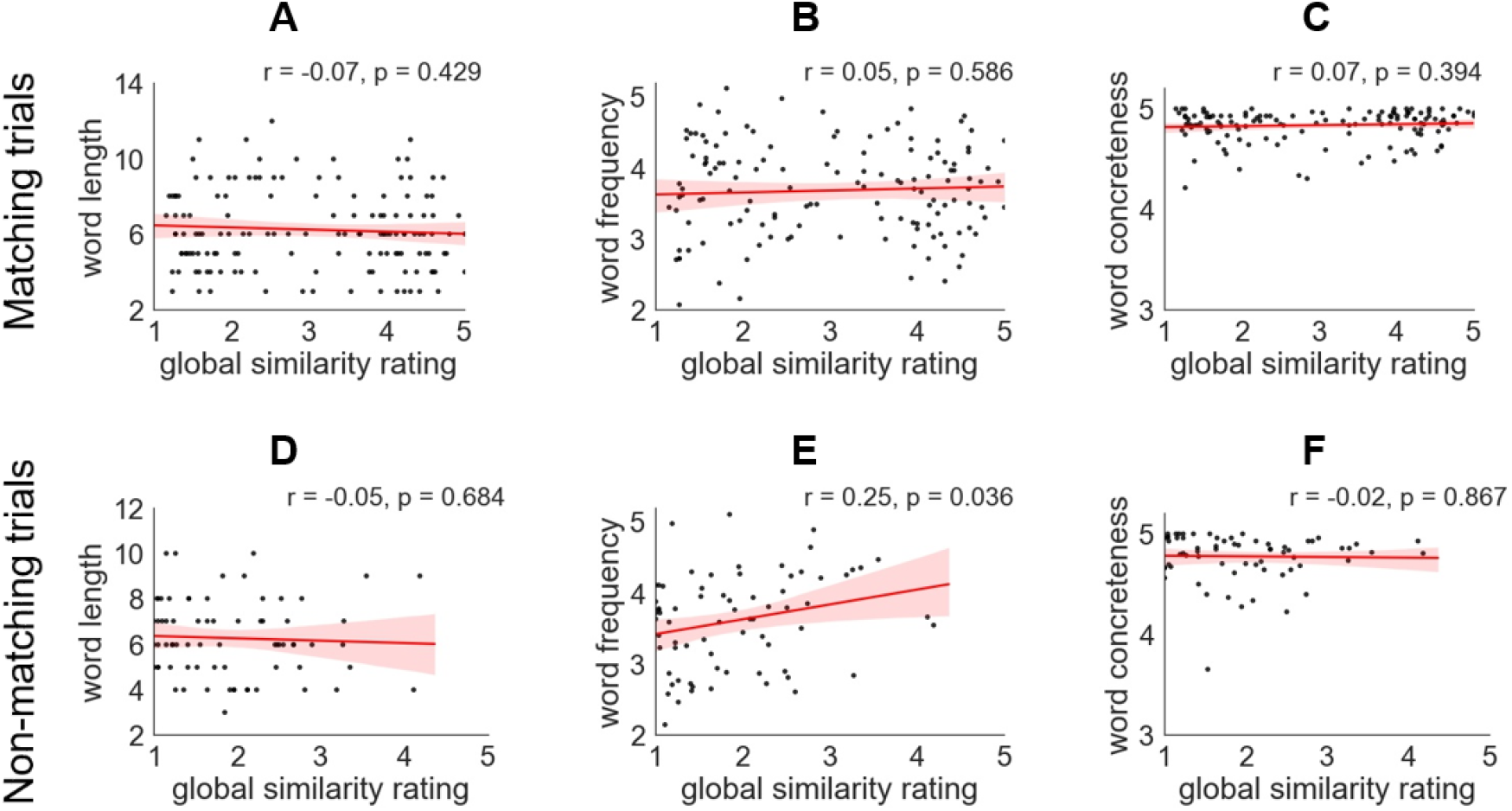
A, B and C show the correlations between global similarity ratings and word length, frequency and concreteness of the target word for matching trials. D, E and F show the correlations for non-matching trials.

**Figure S2.**
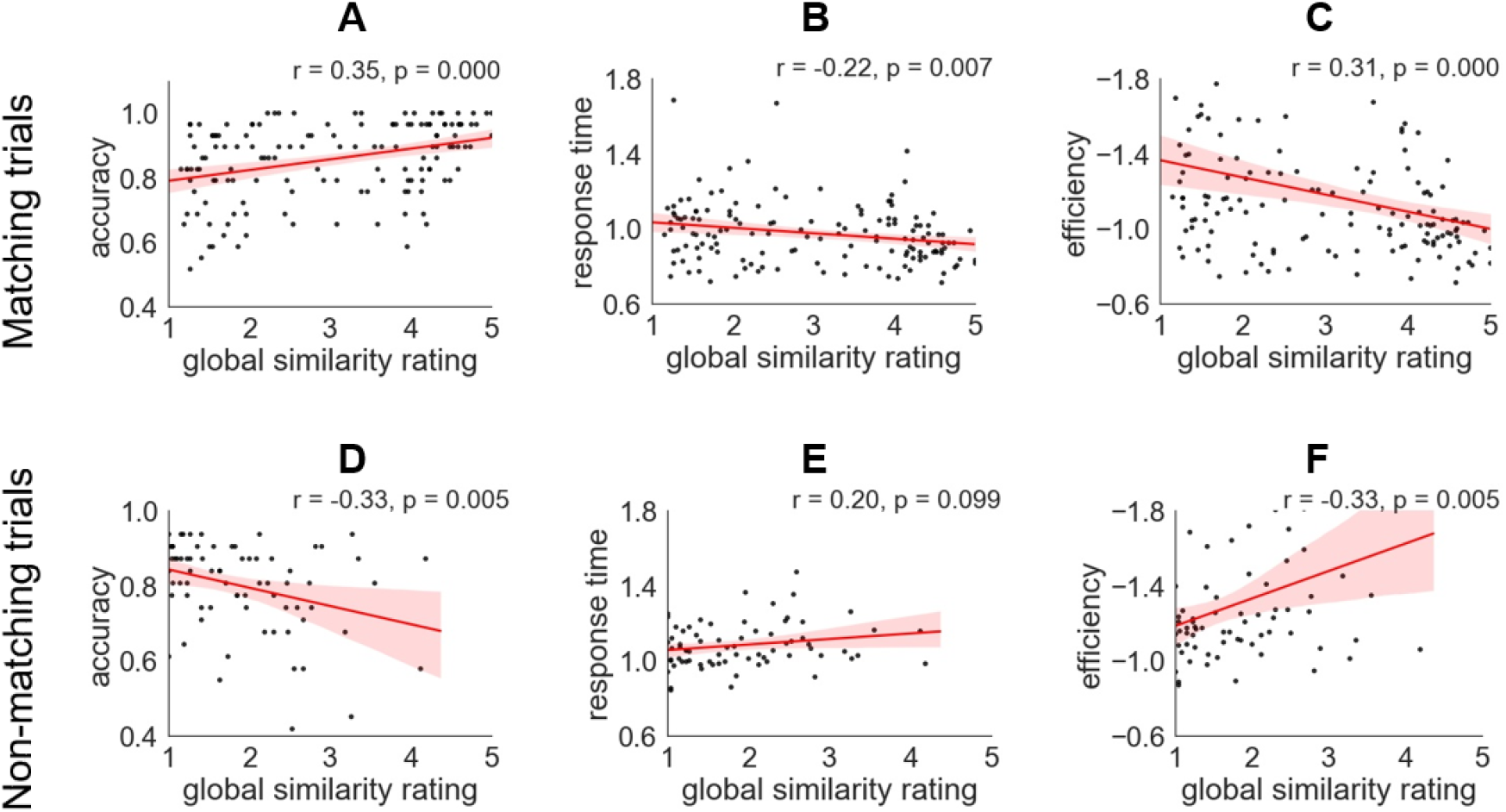
A, B and C show correlations between global similarity ratings and average behavioural performance across 30 participants for the matching trials, examining accuracy, response time and efficiency. D, E and F show the correlations for the non-matching trials. (Each trial is shown as a data point).

**Figure S3.**
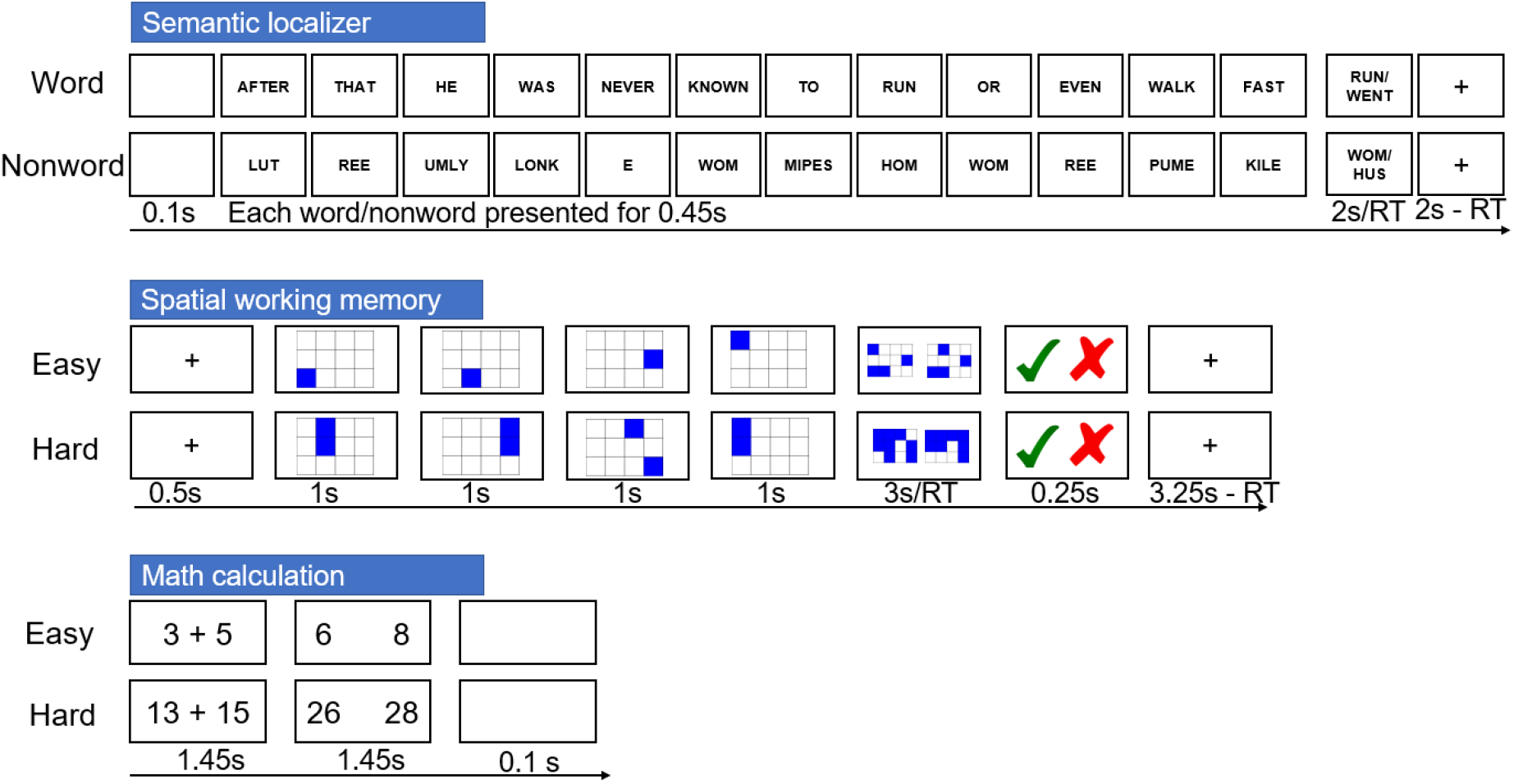
Illustration of localizer tasks.

**Figure S4.**
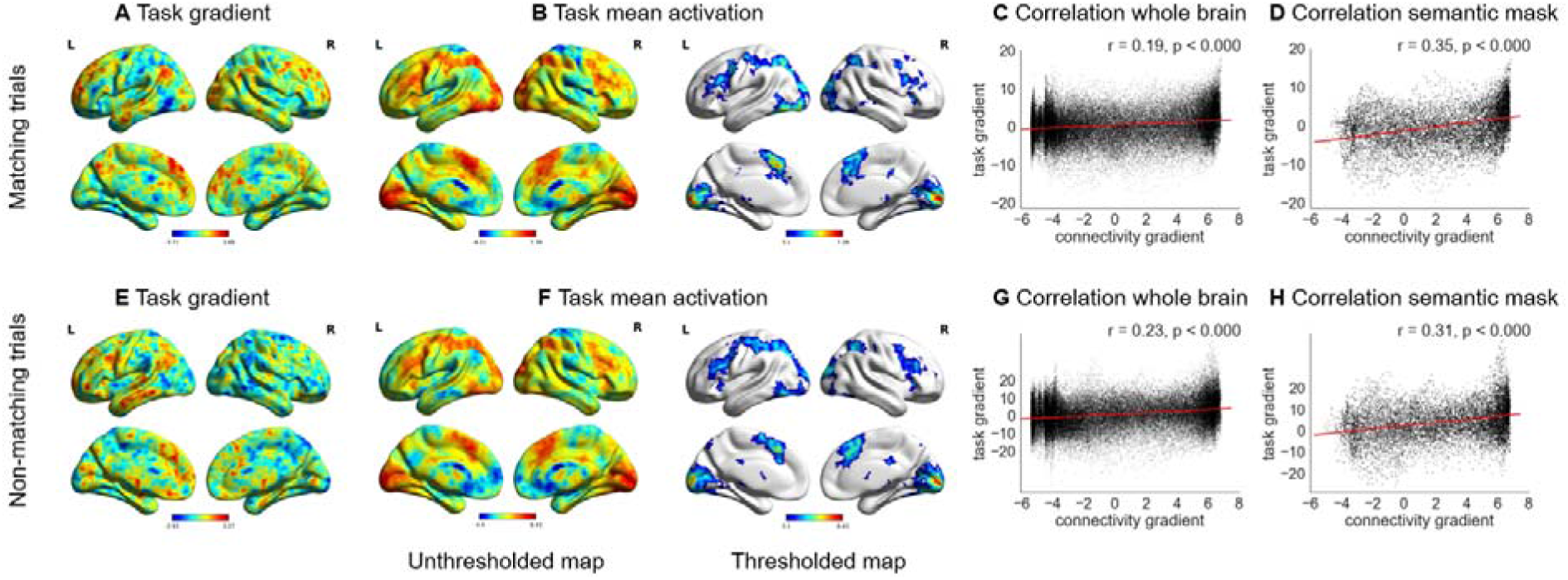
A - Unthresholded map of the task gradient for matching trials: i.e. parametric manipulation of global semantic similarity. Warm colours = positively correlated activity [stronger response when task-irrelevant semantic features were shared between probe and target]. Cool colours = negatively correlated activity [stronger response when only the goal feature was shared between probe and target]. B - Unthresholded and thresholded (FWE-corrected, z = 3.1, p < 0.05) mean effect of the feature matching task for matching trials, which corresponds to the intercept in the parametric analysis. C and D – Correlations between the task gradient for matching trials and the connectivity gradient ^1^ across the whole brain and within the semantic mask defined using Neurosynth. E - Unthresholded map of the task gradient for non-matching trials: i.e. parametric manipulation of global semantic similarity. Warm colours = positively correlated activity [stronger response when task-irrelevant semantic features were shared between probe and target]. Cool colours = negatively correlated activity [stronger response when no features were shared between probe and target]. F - Unthresholded and thresholded (FWE-corrected, z = 3.1, p < 0.05) mean effect of the feature matching task for non-matching trials, which corresponds to the intercept in the parametric analysis. G and H – Correlations between the task gradient for non-matching trials and the connectivity gradient ^1^ across the whole brain and within the semantic mask defined using Neurosynth.

**Figure S5.**
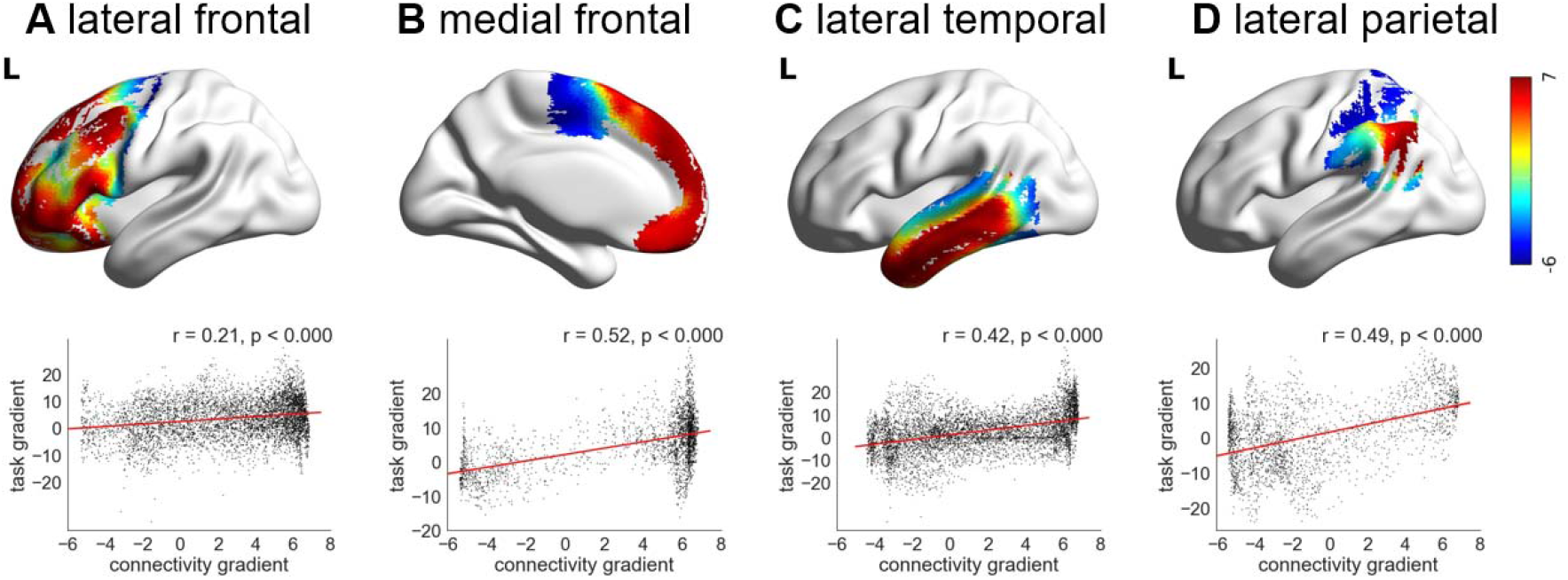
A, B, C and D show the correlations between the task gradient for non-matching trials and the connectivity gradient^1^ in lateral frontal, medial frontal, lateral temporal and lateral parietal cortex, respectively. The colour represents each voxel’s value on the connectivity gradient defined by Margulies et al.^1^. See main text for equivalent correlations for the matching trials.

**Figure S6.**
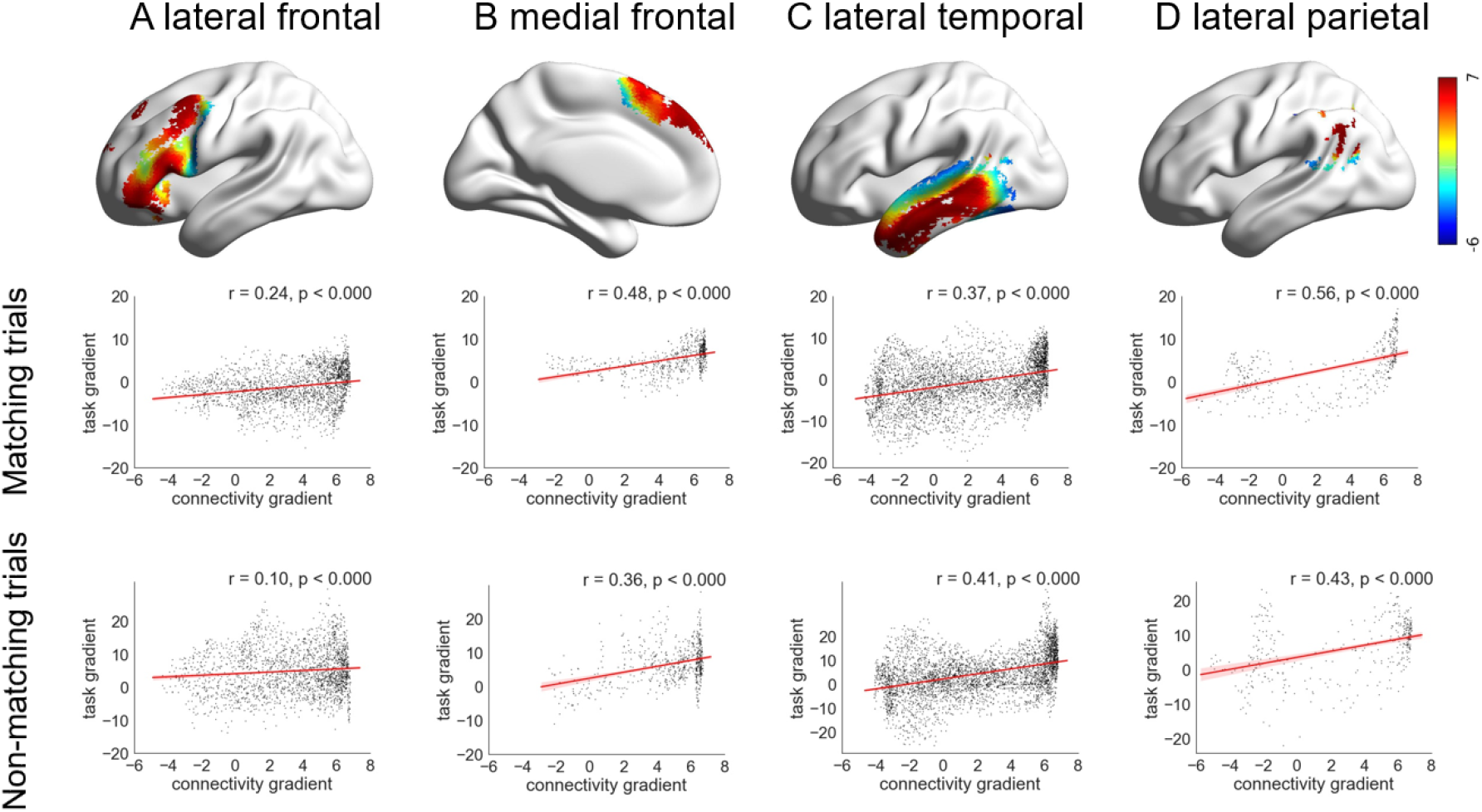
Top panel: A, B, C and D show anatomical regions in lateral frontal, medial frontal, lateral temporal and lateral parietal zones, only including voxels identified as relevant to semantic processing using Neurosynth^18^. The colour represents the connectivity gradient value from Margulies et al.^1^, with red towards the heteromodal end and blue towards the unimodal end of the connectivity gradient. Middle panel: The correlations between the task gradient for matching trials and the connectivity gradient^1^ in each anatomical region shown in the top panel. Bottom panel: The correlations between the task gradient for non-matching trials and the connectivity gradient ^1^ in each anatomical region shown in the top panel.

**Figure S7.**
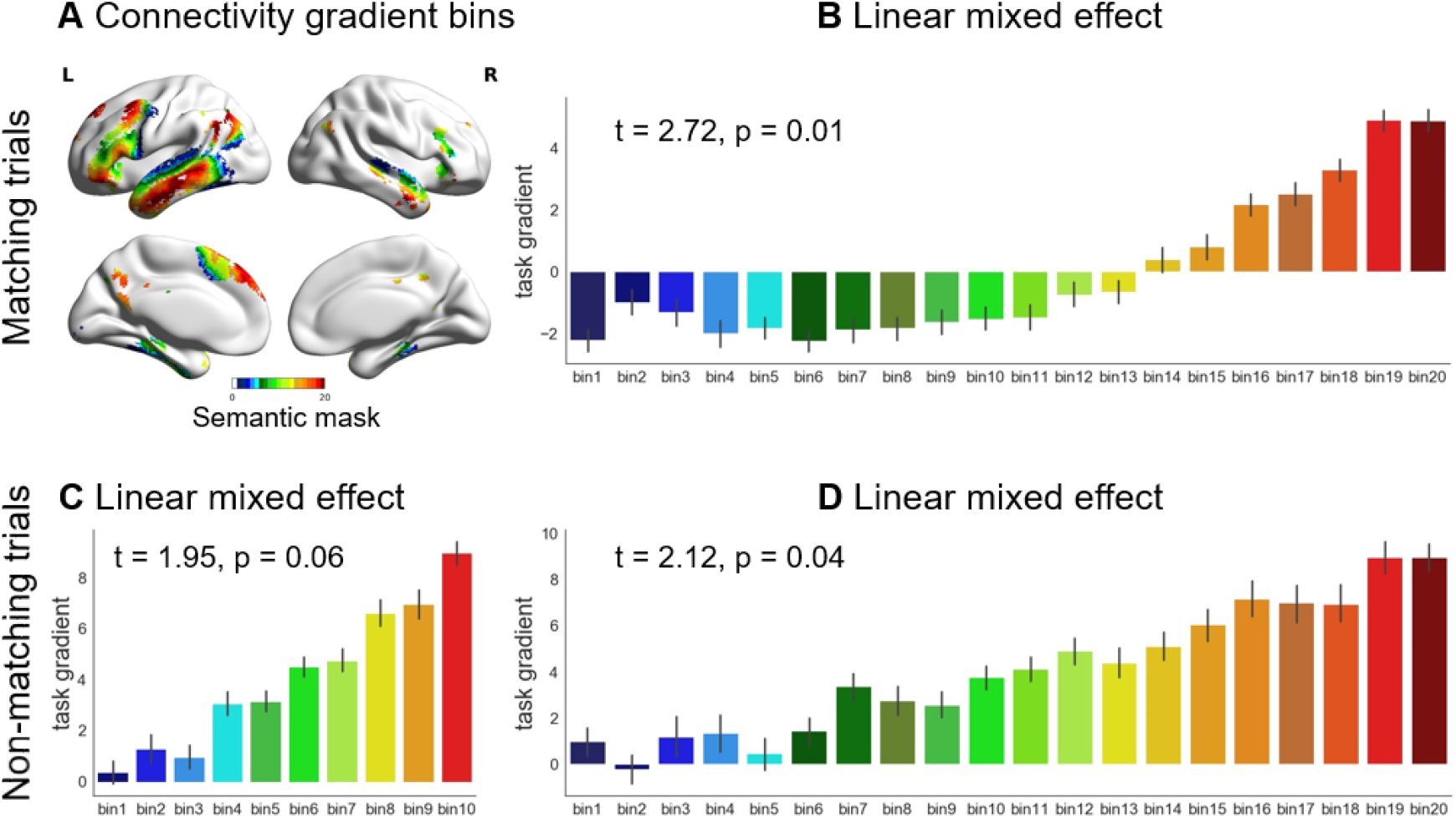
A - The connectivity gradient ^1^ within a semantic mask defined using Neurosynth, divided into 20 bins according to gradient value: bin 1 is located towards the unimodal end, while bin 20 is at the heteromodal end of the principal gradient. B - The effect of global semantic similarity (i.e. the task gradient) for matching trials within each bin of the connectivity gradient within the semantic mask, showing that the response to the task changes in an orderly way. C and D - The effect of the task gradient for non-matching trials in each bin of the connectivity gradient within the semantic mask. This is divided into 10 bins and 20 bins, respectively, showing that the response to the task changes in an orderly way.

**Table S1:** Relationship between connectivity and task gradient for matching trials examined using linear mixed effects modelling using 10 bins. Table of fixed effects.

| Parameter | Estimate | Standard Error | t | p |
| --- | --- | --- | --- | --- |
| Intercept | -3.69 | 2.24 | -1.65 | .11 |
| Connectivity gradient linear effect | 0.55 | 0.23 | 2.36 | .02 |

**Table S2:** Relationship between connectivity and task gradient for matching trials examined using linear mixed effects modelling using 10 bins. Table of random effects.

| Groups | Variance | SD |
| --- | --- | --- |
| Subject intercept | 87.34 | 9.35 |
| Subject slope | 0.25 | 0.50 |
| Run intercept | 1.32 | 1.15 |
| Residual | 443.70 | 21.06 |

**Table S3:** Relationship between connectivity and task gradient for matching trials examined using linear mixed effects modelling using 20 bins. Table of fixed effects.

| Parameter | Estimate | Standard Error | t | p |
| --- | --- | --- | --- | --- |
| Intercept | -3.60 | 2.03 | -1.77 | 0.09 |
| Connectivity gradient linear effect | 0.28 | 0.10 | 2.72 | 0.01 |

**Table S4:** Relationship between connectivity and task gradient for matching trials examined using linear mixed effects modelling using 20 bins. Table of random effects.

| Groups | Variance | SD |
| --- | --- | --- |
| Subject intercept | 82.64 | 9.09 |
| Subject slope | 0.14 | 0.38 |
| Run intercept | 2.09 | 1.45 |
| Residual | 447.27 | 21.15 |

Note: The random effect of run was not included as this model failed to converge.

**Table S5:**
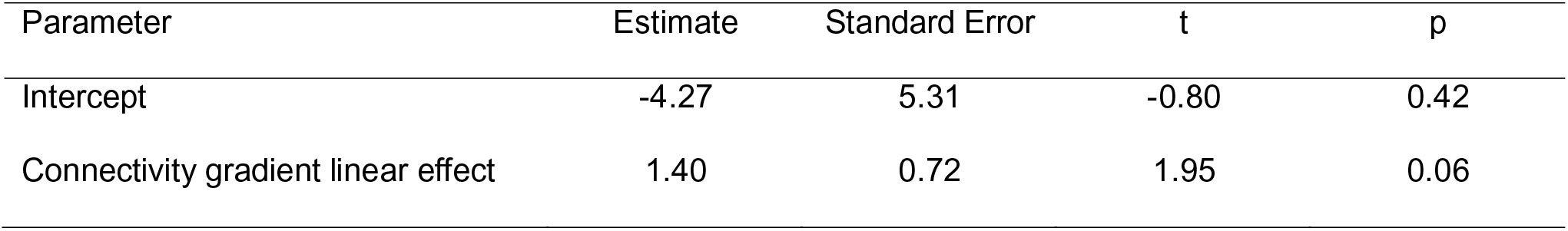
Relationship between connectivity and task gradient examined using linear mixed effects modelling using 10 bins for the non-matching trials. Table of fixed effects.

**Table S6:**
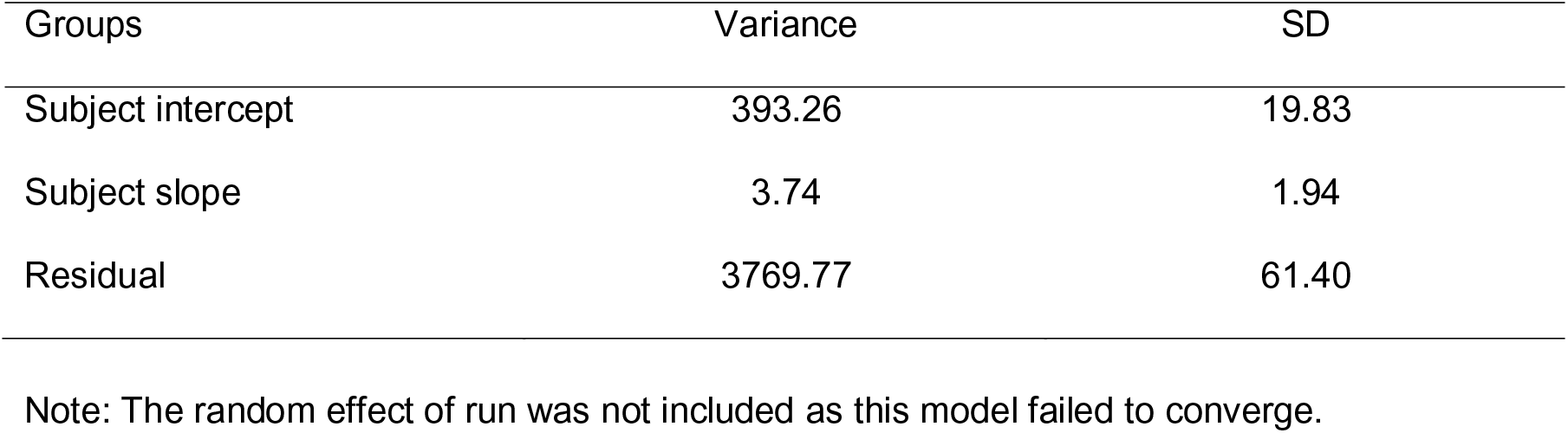
Relationship between connectivity and task gradient examined using linear mixed effects modelling using 10 bins for the non-matching trials. Table of random effects.

**Table S7:** Relationship between connectivity and task gradient examined using linear mixed effects modelling using 20 bins for the non-matching trials. Table of fixed effects.

| Parameter | Estimate | Standard Error | t | p |
| --- | --- | --- | --- | --- |
| Intercept | -4.09 | 5.05 | -0.81 | 0.42 |
| Connectivity gradient linear effect | 0.73 | 0.34 | 2.12 | 0.04 |

**Table S8:** Relationship between connectivity and task gradient examined using linear mixed effects modelling using 20 bins for the non-matching trials. Table of random effects.

| Groups | Variance | SD |
| --- | --- | --- |
| Subject intercept | 501.27 | 22.39 |
| Subject slope | 2.04 | 1.43 |
| Run intercept | 6.63 | 2.58 |
| Residual | 3813.83 | 61.76 |

